# The language network reliably ‘tracks’ naturalistic meaningful non-verbal stimuli

**DOI:** 10.1101/2022.04.24.489316

**Authors:** Yotaro Sueoka, Alexander Paunov, Anna Ivanova, Alyx Tanner, Idan A. Blank, Evelina Fedorenko

## Abstract

The language network, comprised of brain regions in the left frontal and temporal cortex, responds robustly and reliably during language comprehension but shows little or no response during many non-linguistic cognitive tasks (e.g., Fedorenko & Blank, 2020). However, one domain whose relationship with language remains debated is semantics—our conceptual knowledge of the world. Given that the language network responds strongly to meaningful linguistic stimuli, could some of this response be driven by the presence of rich conceptual representations encoded in linguistic inputs? In this study, we used a naturalistic cognition paradigm to test whether the cognitive and neural resources that are responsible for language processing are also recruited for processing semantically rich non-verbal stimuli. To do so, we measured BOLD responses to a set of ∼5-minute-long video and audio clips that consisted of meaningful event sequences but did not contain any linguistic content. We then used the inter-subject correlation (ISC) approach (Hasson et al., 2004) to examine the extent to which the language network ‘tracks’ these stimuli, i.e. exhibits stimulus-related variation. Across all the regions of the language network, non-verbal meaningful stimuli elicited reliable ISCs. These ISCs were higher than the ISCs elicited by semantically impoverished non-verbal stimuli (e.g., a music clip), but substantially lower than the ISCs elicited by linguistic stimuli. Our results complement earlier findings from controlled experiments (e.g., Ivanova et al., 2021) in providing further evidence that the language network shows some sensitivity to semantic content in non-verbal stimuli.

## 1. Introduction

A set of brain regions in the left frontal and temporal cortex in the human brain respond robustly and reliably during language comprehension, both auditory and visual (e.g., Fedorenko et al., 2010; Silbert et al., 2014; Vagharchakian et al., 2012). In brain imaging studies, these regions exhibit highly selective responses to language processing over diverse cognitive tasks, including those that bear parallels to language and have long been argued to share resources with language. For example, language-responsive regions show little or no response to arithmetic (e.g., Fedorenko et al., 2011; Monti et al., 2012; Amalric & Dehaene, 2019), music (e.g., Fedorenko et al., 2011; Rogalsky et al., 2011; Chen et al., 2023), logic (e.g., Monti et al., 2009), executive control (e.g., Fedorenko et al., 2011; Mineroff, Blank et al., 2018), action/gesture observation (e.g., Pritchett et al., 2018; Jouravlev et al., 2019), Theory of Mind (e.g., Deen et al., 2015; Paunov, 2018; Paunov et al., 2022; Shain, Paunov, Chen et al., 2022), and even the processing of computer languages (e.g., Ivanova et al., 2020; Liu et al., 2020). Evidence from patients with aphasia converges with these findings, showing that the loss of language production and comprehension abilities can leave other cognitive abilities unimpaired (e.g., Lecours & Joanette, 1980; Varley & Siegal, 2000; Varley et al., 2001, 2005; Apperly et al., 2006; Bek et al., 2010; Willems et al., 2011; see Fedorenko & Varley, 2016, for a review).

However, one domain whose relationship with language remains debated is semantic knowledge—generalized, abstract conceptual information about objects/entities, actions, and events (Warrington, 1975; Patterson et al, 2007; Lambon Ralph et al, 2017). On the one hand, language understanding is inextricably linked to semantics. Whether producing or comprehending a sentence, accessing the meaning of words and constructions is an essential component of linguistic processing (e.g., Katz & Fodor, 1963; Pustejovsky, 2005; Goldberg, 2010). On the other hand, semantic knowledge is not restricted to language alone: non-verbal stimuli, such as pictures and sounds, can also evoke rich conceptual representations. Here, we ask: is the language network—or some of its components—sensitive to semantic knowledge regardless of whether or not it is delivered linguistically?

Previous work provides conflicting evidence about the role of the language network in non-verbal semantic processing. A number of brain imaging investigations have reported overlapping activations in the lateral frontal and temporal brain areas for the processing of words and object pictures (e.g., Visser et al., 2012; Devereaux et al., 2013; Handjaras et al., 2017), words and actions (Wurm & Caramazza, 2019), or sentences and event pictures (e.g., Ivanova et al., 2021). However, neuropsychological evidence from individuals with aphasia indicates that linguistic and semantic processing are neurally dissociable, such that semantic processing can appear intact in the presence of severe linguistic deficits (e.g., Whitehouse et al., 1978; Chertkow et al., 1997; Saygin et al., 2004; Colvin et al., 2019; Ivanova et al., 2021; Warren & Dickey, 2021).

Most past brain imaging studies that have reported overlap between responses to verbal and semantically meaningful non-verbal (typically, pictorial) stimuli have relied on traditional task-based paradigms. In such paradigms, participants read words or view pictures and perform a semantic task on those words/pictures. One possible explanation of these results is that language regions respond during the processing of pictures because participants activate the corresponding verbal labels as they come up with the response to the (often verbally presented) task prompt (e.g., activating the word “hammer” when processing a picture of a hammer and deciding its semantic category; Devereux et al., 2013). Activating the label may help hold the concept in working memory or retrieve relevant semantic knowledge but not be *essential* to semantic processing per se (see also Benn, Ivanova et al., 2023 for discussion).

In this study, we aimed to determine whether the language network responds to non-linguistic semantically meaningful stimuli in the absence of an explicit task. To do so, we turned to naturalistic stimuli (Hasson et al., 2018) and leveraged the inter-subject correlation (ISC) approach (Hasson et al., 2004, 2008). In this approach, the similarity of neural fluctuations during some naturalistic stimulus is computed across participants, and the degree of inter-subject similarity is taken to reflect the degree to which a given voxel or brain region responds to features of the relevant stimulus (or ‘tracks’ the stimulus).

Our critical stimuli consisted of ∼5-minute-long video and audio clips that were rich in semantic content but did not contain any language. The key operational criteria we used to select stimuli with rich semantic content included the presence of (a) objects/entities/people, actions, and events (e.g., “bird” depicted visually; “chasing” depicted visually; “brushing teeth” conveyed auditorily with a prototypical sound, etc.), and (b) a storyline, such that the sequences of events are sensible (e.g., a man falls asleep at church and rolls off the pew, or a sound of an alarm clock is followed by a sound of a yawn) and interpretation would become more difficult if the event order was shuffled. The latter criterion is another feature that distinguishes our study from most prior neuroimaging studies examining verbal/non-verbal semantic processing (cf. Thierry & Price, 2006; Baldassano et al., 2018), where participants typically view a series of discrete temporally unconnected stimuli. Naturalistic meaningful stimuli that unfold over time in a task-free setting provide a good approximation to many real-life scenarios and can yield important insights about the role of the language network in semantic processing. In addition, a meaningful storyline helps maintain the participants’ attention even in the absence of an explicit task. For comparison, we also included naturalistic linguistic stimuli, which have been previously shown to elicit high ISCs in the language network (e.g., Wilson et al., 2007; Lerner et al., 2011; Blank & Fedorenko, 2017; Fedorenko & Blank, 2020; Paunov et al., 2022), and non-verbal semantically impoverished stimuli, like music, which serve as negative controls. For simplicity, we refer to semantically rich stimuli as “meaningful”, or +M and semantically impoverished stimuli as “non-meaningful”, or -M.

To foreshadow the key results, we found that the language network shows significantly higher ISCs during the processing of non-verbal meaningful stimuli compared to non-verbal non-meaningful stimuli. These ISCs were nonetheless lower than those elicited during the processing of linguistic stimuli, in line with abundant evidence that linguistic stimuli are the preferred stimulus class for the language network. Our findings suggest that the language network is recruited for the processing of semantically rich inputs even in the absence of language.

## 2. Materials and Methods

### 2.1. General Approach

To investigate whether the language brain regions track semantic content in non-verbal stimuli, we combined two methodologies commonly used in neuroimaging research (similar to e.g., Blank & Fedorenko, 2017): functional localization (e.g., Brett et al., 2002; Saxe et al., 2006; Fedorenko et al., 2010) and rich naturalistic stimuli analyzed with the inter-subject correlation (ISC) approach (Hasson et al., 2004, 2008). In this section, we motivate these methodological choices.

First, to account for inter-individual variability in the location of the language network (e.g., Amunts et al., 1999; Tomaiuolo et al., 1999; Fedorenko et al., 2010; Fedorenko & Blank, 2020; Lipkin et al., 2022), we defined language regions in each participant individually using a well-established functional localizer task (Fedorenko et al., 2010). This approach yields greater sensitivity, functional resolution, and interpretability compared to the traditional group-averaging approach, where functional correspondence across participants is assumed to hold voxel-wise (e.g., Nieto-Castañón & Fedorenko, 2012; Fedorenko, 2021).

And second, naturalistic stimuli are more ecologically valid and thus more likely to elicit neural responses that reflect how the target brain region or network is recruited during real-life cognitive processing. However, the less constrained nature of the stimuli poses a challenge when interpreting the observed blood oxygen-level dependent (BOLD) signal, namely that there are no well-defined conditions within a stimulus from which to construct model-based predictors of neural activity over time. Here, we rely on inter-subject correlations (ISCs), a method introduced by Hasson et al. (2004) to tackle this issue. The reasoning behind the ISC approach is as follows: if a voxel or a brain region tracks certain features of a stimulus, then the BOLD signal should be time-locked to the appearance of those features in the stimulus, which would be reflected in similar patterns of fluctuations across participants (i.e., reliable ISCs). Thus, the average timecourse across participants serves as the “model” for a given left-out participant.

In the next sections, we detail each analysis method and describe other aspects of the study’s design and procedure. Data and code used for this study is available on OSF (https://osf.io/t8sz5/?view_only=d5081a3bf15942c38c7d48f093eac16c).

### 2.2. Participants

Forty-nine native English speakers between the ages of 19 and 48, recruited from MIT and the surrounding community, were paid for participation. Two participants were removed from the analysis due to poor quality of the functional localizer data, with the exclusion criterion defined as fewer than 100 suprathreshold voxels (at the p *<* 0.001 uncorrected whole-brain threshold) across the language network’s ‘parcels’ (**Figure 1B**). Of the remaining 47 participants (mean age = 24.4, SD = 5.08; 29 females), seven participants were left-handed (based on the Edinburgh Handedness Inventory; Oldfield, 1971), but all of those left-handed participants had a left-lateralized language network (for motivation to include left-handers in cognitive neuroscience research, see Willems et al., 2014). All participants gave informed written consent in accordance with the requirements of MIT’s Committee on the Use of Humans as Experimental Subjects (COUHES).

**Figure 1.**
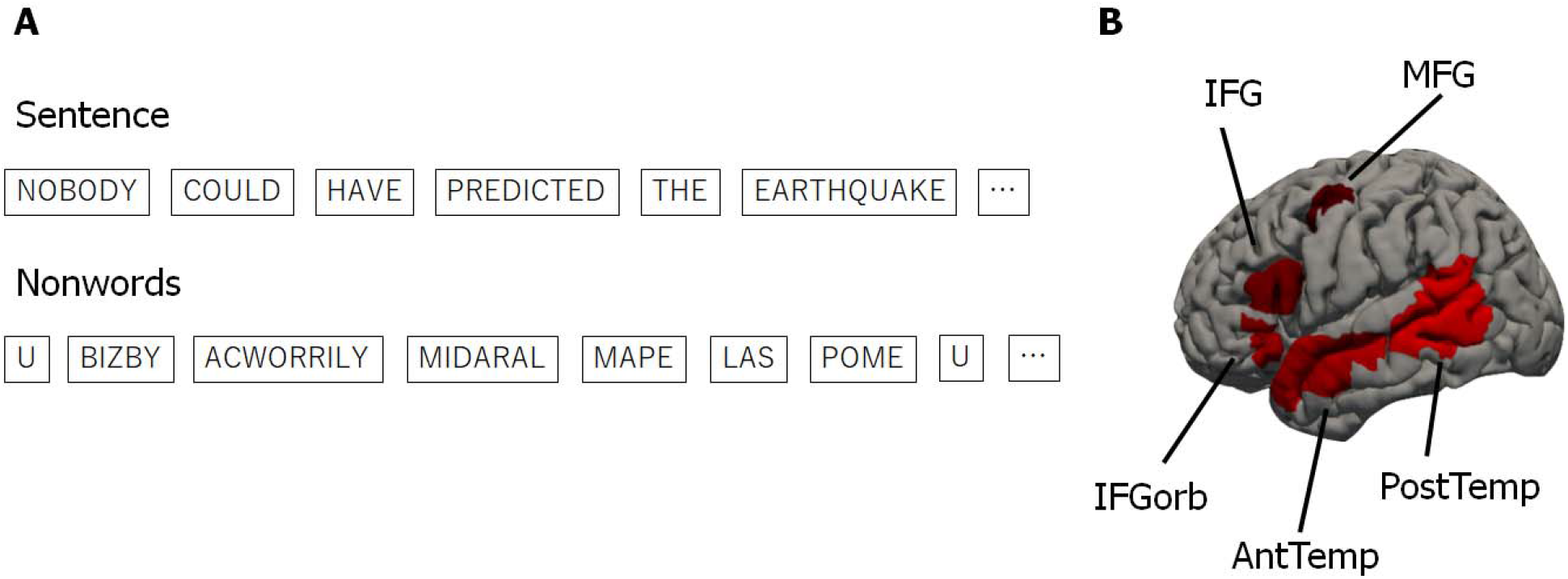
Language Localizer Task. **A.** An example sentence / nonword sequence presented during the language localizer task. Words / nonwords were presented one at a time for 450ms each. **B.** Five language parcels that were used to define the language network for each individual. IFGorb = inferior frontal gyrus, orbital portion; IFG = inferior frontal gyrus; MFG = middle frontal gyrus; AntTemp = anterior temporal lobe; PostTemp = posterior temporal lobe. Each participant’s functional ROIs were defined as the top 10% of most language-responsive voxels within each parcel.

### 2.3. Design and procedure

Each participant completed a language localizer and a subset of the naturalistic stimuli from the critical experiment. As shown in **Table 1**, the majority of the participants (n=27, 57%) watched/heard 8 of the 10 stimuli, one participant—all 10 stimuli, and the remaining 19 participants saw between 1 and 7 stimuli due to scan duration constraints (the differences in the number of stimuli are accounted for in the analyses, as described below). Each stimulus was presented to between 14 and 34 participants, and each +M stimulus was presented to at least 29 participants (**Table 2**). 40/47 participants completed the language localizer in the same session as the critical experiment, the remaining 7 participants completed it in an earlier session (see Mahowald & Fedorenko, 2016 and Lipkin et al., in prep. for evidence of high across-session replicability; also Braga et al., 2020). Some participants completed additional experiments for unrelated studies.

**Table 1.** Number of participants who watched / heard each subset of naturalistic conditions.

| Number of<br>Stimuli | Number of<br>Participants |
| --- | --- |
| 1 | 7 |
| 2 | 6 |
| 3 | 1 |
| 4 | 1 |
| 6 | 1 |
| 7 | 3 |
| 8 | 27 |
| 10 | 1 |

**Table 2.**
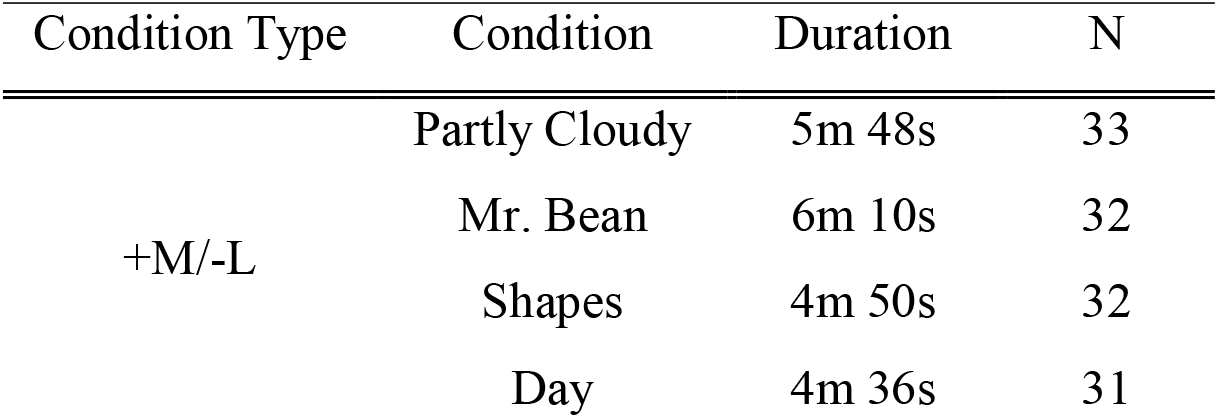
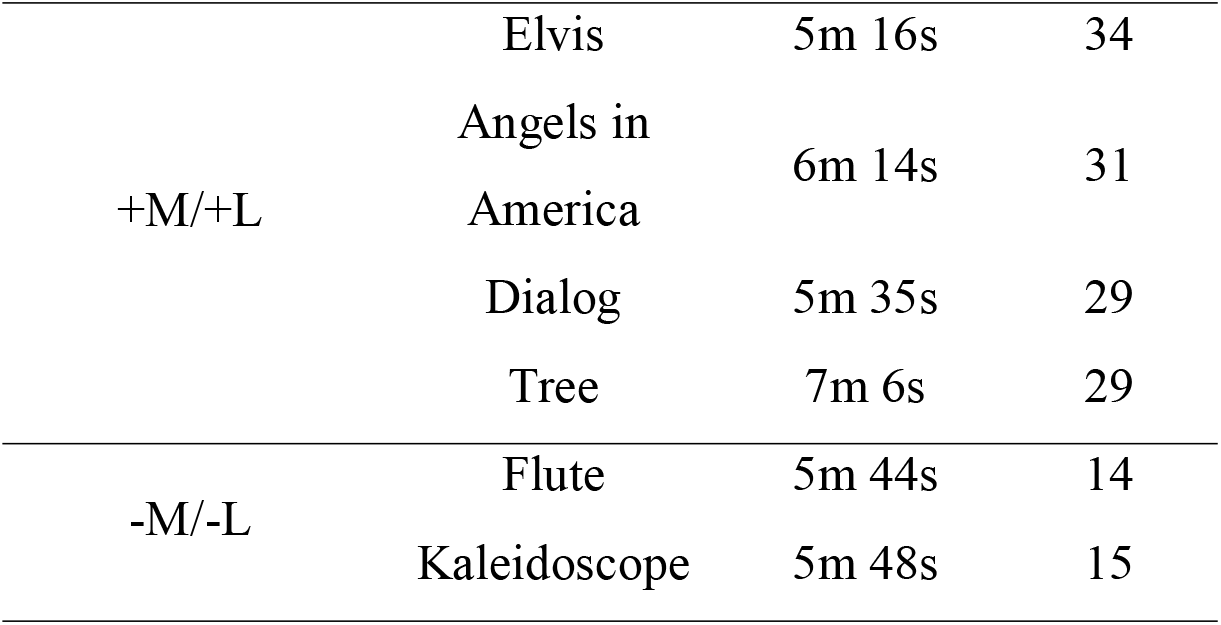
Details of the naturalistic conditions. Each condition belonged to one of the three condition types. The duration of each stimuli, as well as number of participants who viewed / listened to each stimuli (N) are shown. Durations include 16s-long fixations at the beginning and the end of the scan (32s total). +M/-L = +Meaning/-Language; +M/+L = +Meaning/+Language; -M/-L = -Meaning/-Language.

#### 2.3.1. Language localizer

Participants read sentences (e.g., NOBODY COULD HAVE PREDICTED THE EARTHQUAKE IN THIS PART OF THE COUNTRY) and lists of unconnected, pronounceable nonwords (e.g., U BIZBY ACWORRILY MIDARAL MAPE LAS POME U TRINT WEPS WIBRON PUZ) in a blocked design (**Figure 1A**). Each stimulus consisted of twelve words/nonwords. For details of how the language materials were constructed, see Fedorenko et al. (2010). The materials are available at https://evlab.mit.edu/funcloc/. The sentences > nonword-lists contrast has been previously shown to reliably activate left-lateralized frontal and temporal language processing regions and to be robust to changes in the materials, task, and modality of presentation (Fedorenko et al., 2010; Mahowald & Fedorenko, 2016; Scott et al., 2017). Stimuli were presented in the center of the screen, one word/nonword at a time, at the rate of 450 ms per word/nonword. Each stimulus was preceded by a 100 ms blank screen and followed by a 400 ms screen showing a picture of a finger pressing a button, and a blank screen for another 100 ms, for a total trial duration of 6 s. Participants were asked to press a button whenever they saw the picture of a finger pressing a button. This task was included to help participants stay alert and awake. Condition order was counterbalanced across runs. Experimental blocks lasted 18 s (with 3 trials per block), and fixation blocks lasted 14 s. Each run (consisting of 5 fixation blocks and 16 experimental blocks) lasted 358 s. Each participant completed 2 runs.

#### 2.3.2. Critical experiment

Participants passively watched or listened to a set of naturalistic stimuli (**Figure 2A**). Auditory stimuli were presented over scanner-safe Sensimetrics headphones. Each of 10 stimuli (conditions) belonged to one of 3 condition types: +Meaning/-Language, +Meaning/+Language, and -Meaning/-Language (**Figure 2**). The stimuli within each condition type were chosen to vary in their presentation modalities (visual/auditory) and styles (e.g., for visual stimuli: live action/animated movie/abstract shapes) to ensure generalizability within a condition type. The four stimuli from the critical condition were rich in semantic content but did not contain any language (+Meaning/-Language; +M/-L): i) an animated short film (“Partly Cloudy” by Pixar), ii) a clip from a live action film (“Mr. Bean”, https://www.youtube.com/watch?v=bhg-ZZ6WA), iii) a custom-created Heider and Simmel-style animation (Heider & Simmel, 1944) consisting of simple geometric shapes moving in ways as to suggest intentional interactions designed to tell a story (e.g., a shape gets locked up inside a space, another shape goes on a quest to get help to release it, etc.), denoted as “Shapes” hereafter, and iv) a custom-created audio ‘story’ consisting of a series of sounds that represents common events in a typical morning, denoted as “Day” hereafter. Four stimuli from the positive control condition used semantically rich verbal materials (+Meaning/+Language; +M/+L): i) a story (“Elvis” from the Natural Stories corpus; Futrell et al., 2020), ii) an audio play (a segment from an HBO miniseries, “Angels in America”, from Chapter 1: https://www.imdb.com/title/tt0318997/), iii) a naturalistic dialog—a casual unscripted conversation between two female friends, denoted as “Dialog” hereafter, and iv) an expository text (a text about trees adapted from Wikipedia; https://en.wikipedia.org/wiki/Tree), denoted as “Tree” hereafter. The two stimuli from the negative control condition lacked both semantic and verbal content (-Meaning/-Language; -M/-L): i) an audio clip of Paganini Caprice No. 24 arranged for solo flute, denoted as “Flute” hereafter, and ii) a video consisting of continuously changing kaleidoscope images, denoted as “Kaleidoscope” hereafter. Each stimulus, lasting ∼5-7 minutes (see **Table 2** for detailed timing information), was preceded and followed by 16 s of fixation. All the materials are available on OSF (https://osf.io/t8sz5/?view_only=d5081a3bf15942c38c7d48f093eac16c), except “Partly Cloudy” (for copyright reasons).

**Figure 2.**
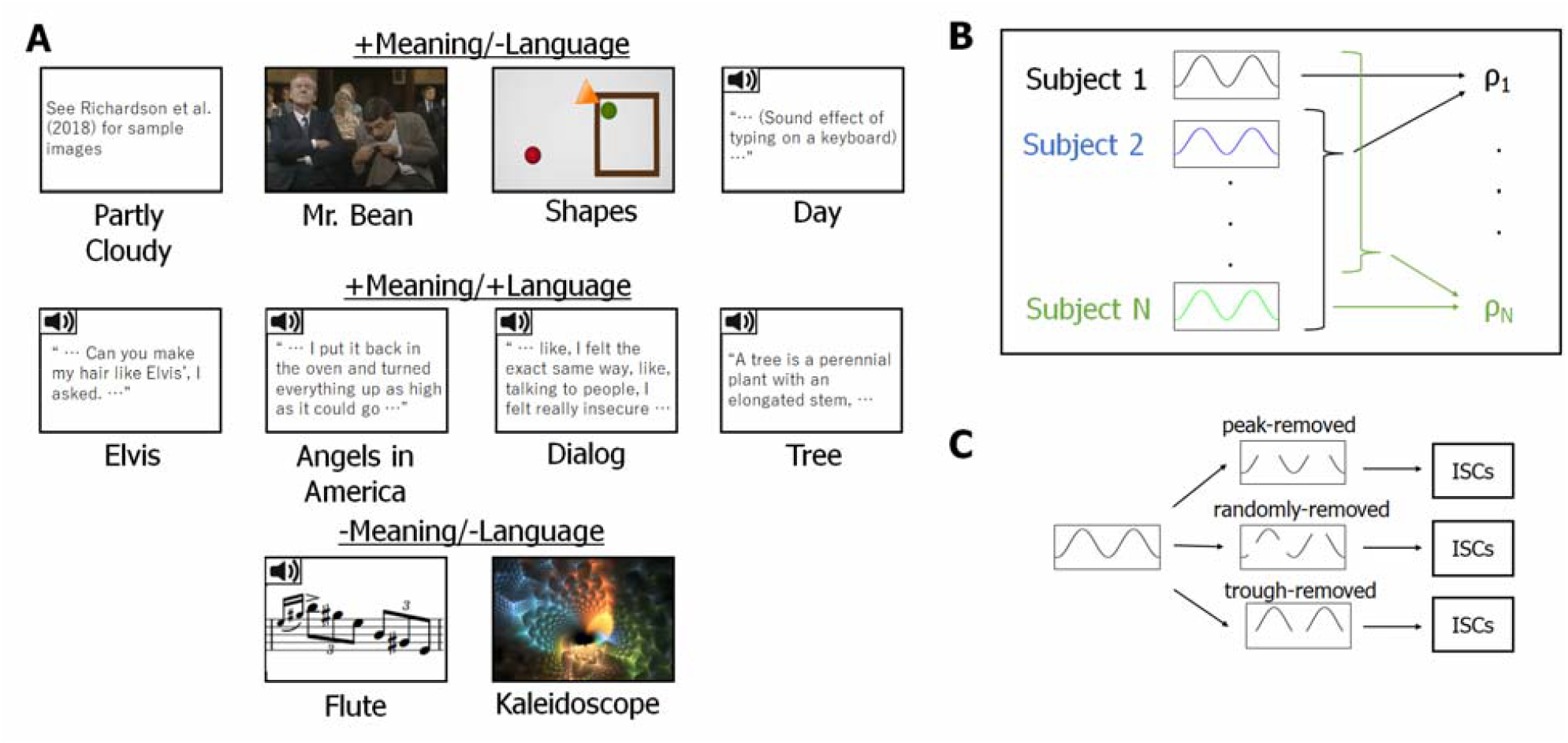
Critical task and schematics of analysis. **A.** Ten conditions used in the critical task. The speaker icons represent auditory stimuli, while others are visual stimuli. **B.** Schematics of ISC calculation. ISCs were computed by correlating each participant’s BOLD time series for a particular condition in a particular fROI with the average of the other participants’ BOLD time series for that condition in that fROI and Fisher-transforming the correlation coefficients. These ISCs were used for analysis with linear mixed effect models. **C.** Schematics for the peak-removal analysis. From each time course, peak, trough, or random time points were removed. These trimmed time courses were then used to compute the ISCs as shown in **B**.

#### 2.3.3. Resting state scan

Ten participants additionally performed a resting state scan, used in a control analysis. Participants were instructed to close their eyes and let their mind wander but to remain awake while resting in the scanner for 5 min (the scanner lights were dimmed, and the projector was turned off).

### 2.4. Data acquisition and preprocessing

#### 2.4.1 Data acquisition

Whole-brain structural and functional data were collected on a whole-body 3 Tesla Siemens Trio scanner with a 32-channel head coil at the Athinoula A. Martinos Imaging Center at the McGovern Institute for Brain Research at MIT. T1-weighted structural images were collected in 176 axial slices with 1 mm isotropic voxels [repetition time (TR) = 2530 ms; echo time (TE) = 3.48 ms]. Functional, blood oxygenation level-dependent (BOLD) data were acquired using an EPI sequence with a 90° flip angle and using GRAPPA with an acceleration factor of 2; the following parameters were used: thirty-one 4-mm-thick near-axial slices acquired in an interleaved order (with 10% distance factor), with an in-plane resolution of 2.1 × 2.1 mm, FoV in the phase encoding (A >> P) direction 200 mm and matrix size 96 × 96 mm, TR = 2000 ms and TE = 30 ms. The first 10 s of each run were excluded to allow for steady-state magnetization.

#### 2.4.2. Spatial preprocessing

Data preprocessing was performed with SPM12 (using default parameters, unless specified otherwise) and custom scripts in MATLAB. Preprocessing of anatomical data included normalization into a common space [Montreal Neurological Institute (MNI) template] and segmentation into probabilistic maps of the gray matter (GM), white matter (WM), and cerebro-spinal fluid (CSF). A GM mask was generated from the GM probability map and resampled to 2 mm isotropic voxels to mask the functional data. The WM and CSF maps were used as described in *Temporal preprocessing* below. Preprocessing of functional data included motion correction (realignment to the mean image using second-degree b-spline interpolation), normalization (estimated for the mean image using trilinear interpolation), resampling into 2 mm isotropic voxels, and smoothing with a 4 mm FWHM Gaussian filter.

#### 2.4.3. Temporal preprocessing

The localizer runs were high-pass filtered at 128 s. Additional preprocessing of data from the naturalistic runs was performed using the CONN toolbox (Whitfield-Gabrieli & Nieto-Castañón, 2012; http://www.nitrc.org/projects/conn) with default parameters, unless specified otherwise. BOLD signal time courses were extracted from white matter and cerebrospinal fluid. Five temporal principal components were obtained from each, as well voxel-wise averages. These were regressed out of each voxel’s time course, along with additional noise regressors, specifically, six motion parameters estimated during offline motion correction (three translations and three rotations) and their first temporal derivatives, and artifact timepoints (based on global signal outliers and motion).

### 2.5. Participant-specific functional localization of language network

#### 2.5.1. Modeling localizer data

For the localizer task, a standard mass univariate analysis was performed in SPM12 whereby a general linear model estimated the effect size of each condition in each experimental run. These effects were each modeled with a boxcar function (representing entire blocks) convolved with the canonical hemodynamic response function. The model also included first-order temporal derivatives of these effects, as well as nuisance regressors representing entire experimental runs, offline-estimated motion parameters, and timepoints classified as outliers (i.e., timepoints where scan-to-scan differences in global BOLD signal were above 5 standard deviations, or where the scan-to-scan motion was above 0.9 mm). The obtained weights were then used to compute the functional contrast of interest: sentences > nonwords.

#### 2.5.2. Defining fROIs

Language functional regions of interest (fROIs) were defined individually for each participant based on functional contrast maps from the localizer experiments (a toolbox for this procedure is available online; https://evlab.mit.edu/funcloc/). These maps were first restricted to include only gray matter voxels by excluding voxels that were more likely to belong to either the WM or the CSF based on SPM’s probabilistic segmentation of the participant’s structural data, as described above.

Then, fROIs in the language network were defined using group-constrained, subject-specific localization (Fedorenko et al., 2010). For each participant, the map of the sentences > nonwords contrast was intersected with binary masks that constrained the participant-specific language network to fall within areas where activations for this contrast are likely across the population. These masks are based on a group-level representation of the contrast obtained from an independent large (n=220) sample of participants. We used five such masks in the left-hemisphere, including regions in the inferior frontal gyrus, in its orbital part, in the middle frontal gyrus, in the anterior temporal lobe, and in the mid-to-posterior temporal lobe. These masks are available online (https://evlab.mit.edu/funcloc/). In each of these five masks, a participant-specific language fROI was defined as the 10% of voxels with the highest *t-*values for the sentences > nonwords contrast (Note that this contrast allows for the inclusion of voxels in a fROI that respond above baseline to the nonwords condition, as long as these responses are weaker than to sentences.).

### 2.6. Critical analysis: ISCs

#### 2.6.1. Computing ISCs

For each participant, BOLD signal time courses recorded during each naturalistic condition were extracted from each voxel beginning 6 s following the onset of the stimulus (to exclude an initial rise in the hemodynamic response relative to fixation, which could increase ISCs). These time courses were temporally z-scored in each voxel and then averaged across voxels within each fROI. Then, for each fROI, participant, and condition we computed an ISC value, namely, Pearson’s moment correlation coefficient between the time course and the corresponding average time course across the remaining participants (**Figure 2B**; Lerner et al., 2011). ISCs were Fisher-transformed before statistical testing to improve normality (Silver & Dunlap, 1987).

We also tested whether the relatively high ISCs in +M/-L conditions could be attributed to specific high-response time points. In particular, specific ‘scenes’ in the stimuli might activate the language network across participants because they elicit some linguistic representations. In other words, the tracking of the +M/-L conditions may be due not to the processing of meaningful non-verbal information but instead driven by a few points where participants actually engage in linguistic processing. To evaluate this possibility, ISCs were recomputed after peak time points (or trough time points, as a control) of the BOLD signal were removed (peak-/ trough-removed ISCs, **Figure 2C**). To define peaks, a mean BOLD signal was constructed for each condition by averaging the time course across participants and fROIs. Peak time points were defined as those with values greater than 1 in the z-normalized average time course. The number of time points removed was as follows: Partly Cloudy, 25 out of 158; Mr. Bean, 25 out of 169; Shapes, 20 out of 129; Day, 19 out of 122. Similarly, trough time points were defined as time points with values lower than -1 in the z-normalized average time course. The number of time points removed was as follows: Partly Cloudy, 25 out of 158; Mr. Bean, 24 out of 169; Shapes, 21 out of 129; Day, 16 out of 122.

#### 2.6.2. Statistical testing

In the critical analyses, we compared ISCs across condition types using linear mixed-effects (LME) models (Barr et al., 2013), implemented in R. ISCs were modeled with condition type as a fixed effect and fROI, participant, and condition (stimulus) as random intercepts. The model was tested for significance in two-tailed hypothesis tests. Mixed-effects modeling is more robust to missing-at-random data (i.e., the number of stimuli per participant, see **Table 1**), as well as to unbalanced designs (i.e., different numbers of conditions per condition type). For all LME analyses, we coded condition type using the dummy coding scheme, with +M/-L condition type serving as a reference level, unless specified otherwise.

We additionally investigated ISCs for each +M/-L condition separately by treating each +M/-L condition as individual condition types, which resulted in six condition types in total (+M/+L, - M/-L, Partly Cloudy, Mr. Bean, Shapes, and Day). Here, the main goal was to compare the ISCs during each condition to those observed during the zero baseline. ISCs were modeled with condition type as a fixed effect and fROI and participant as random intercepts with the fixation baseline as a reference level. Similarly, ISCs during each +M/-L condition were compared to mean ISCs during +M/+L and -M/-L condition types. Here, mean ISCs during +M/+L and -M/-L condition types were used as reference levels. P-values in this analysis are reported following false discovery rate (FDR) correction for multiple (n=3; against zero-baseline, mean +M/+L, and mean -M/-L) comparisons (Benjamini & Yekutieli, 1991).

The above network-level analysis was also performed for each fROI. Here, ISCs were modeled for each fROI separately, with condition type as a fixed effect and participant and condition as random intercepts, and p-values were FDR-corrected for multiple comparisons across fROIs (n=5). To test that each condition was above baseline for each fROI, each +M/-L condition was recoded into individual condition types, and ISCs were modeled with condition type as a fixed effect and participant as random intercepts with the fixation baseline as a reference level. Again, p-values were FDR-corrected for multiple comparisons across fROIs (n=5).

Finally, the peak-removed (and trough-removed as a control) ISCs for each condition were compared against a distribution of ISCs (n=10,000) where n time points in the timecourse were pseudo-randomly removed (n matched the number of peak time points for each condition). These adjusted timecourses were generated in MATLAB 2018a. The peak-removed and trough-removed ISCs were also tested against each other using a two-sample t-test for each condition. For the latter analysis, trough-removed ISCs were adjusted so that the number of timepoints removed (the ones with the lowest values) matched the number of peak timepoints removed for each condition. P-values were FDR-corrected for multiple comparisons across conditions (n=4).

### 2.7. Content Unit (CU) Analysis

To measure the degree of semantic content in a given stimulus, we quantified the number of Content Units (CUs) in each clip. We defined a CU as a unique concept (object, entity, action, property, etc.) referred to in verbal descriptions of each stimulus (Agis et al., 2016; Gallée et al., 2021). To estimate the number of CUs in each clip, we conducted an online behavioral study with an independent pool of participants.

The task was implemented as an online Qualtrics survey that presented a randomly chosen stimulus to each participant. The stimulus was cut into ∼30s subclips and the subclips were presented in chronological order. The splitting of stimuli into subclips was done manually to minimize abrupt cuts that would split natural events in the middle; the same subclips were used for all participants for a given stimulus. Each subclip presentation page (participants could only watch each subclip once) was followed by a text box page, where participants were asked to describe the stimulus they just watched / listened to. The transition to the text box page was enabled five seconds after the clip ended. At the end of the survey, as an attention check, participants were asked two questions about the content of the stimulus they had been presented.

For this experiment, 250 native English speakers between the ages of 19 and 77 were recruited online via the Prolific platform. Three participants were removed for failing attention checks at the end of the survey and providing minimal, low-effort responses. Of the 247 remaining participants, twenty participants per stimulus were randomly selected (for a total of 200 participants; n=20 × 10 stimuli) for further analyses.

Before analyzing the descriptions for the number of content units, the following (content-irrelevant) categories of phrases were manually removed: explicitly self-referential language (“I think”, “I believe”, etc.), meta-descriptive language (“Same as before”, “In this clip”, “the camera”, “screen”, etc.), and language regarding levels of confidence/belief states (“It appears to be”, “seems like”, etc.). The remaining portions of the descriptions were stripped of ‘stop words’ (high-frequency function words) using the nltk package in Python, lemmatized using the spaCy package in Python, and split into individual words. Words that served as subcomponents of proper nouns (e.g., “Mr. Bean”, “Elvis Presley”) were merged and counted as a single concept. The same word that appeared multiple times within the response to a single subclip was treated as different concepts because the same entity could participate in multiple events within the clip. Words that appeared in descriptions of three or more participants for a given subclip were defined as CUs.

To compare CU counts across conditions, CU counts were averaged across participants to obtain a single CU count value per subclip. These CU counts were further averaged across subclips to yield a single CU count value for each condition. To statistically compare CU counts between condition types, we used an LME to model average (across participants) subclip CU counts with condition type as a fixed effect and condition (stimulus) as random intercepts. The relationship between CU counts and the ISCs was evaluated, first, with a Pearson correlation coefficient between the average CU count and the average ISCs for each condition. Second, a model comparison examined whether semantic richness (as approximated by CU counts) explained additional variance in ISCs over condition type. We created two models to predict ISCs of stimuli from the three condition types: one adding normalized CU count (centered around 0) as a fixed effect to the original model and the other – replacing condition type with normalized CU count for the fixed effect. The first model was compared with the original model using the likelihood ratio test. The second model was compared with the original model using AIC (Akaike information criterion) given that the two models were not nested. The above analyses were also performed using pairwise shared CU count, defined as the average number of CUs that were shared between each participant pair.

## 3. Results

### 3.1. The language regions reliably track meaningful non-verbal stimuli

Meaningful non-verbal (+M/-L) conditions elicited ISCs that were significantly higher than the zero baseline (mean ISCs = 0.193, beta = 0.168, SE = 0.0434, p = 0.00151; **Figure 3A**). As a sanity check, we also examined the BOLD signal during the resting state condition to ensure that spurious correlations do not arise from the data acquisition and analysis procedures. The ISCs during rest averaged across fROIs were not greater than 0 (one-tailed t-test, t(9) = -1.53, p = 0.920), suggesting that the ISCs for the other conditions reflect stimulus-related activity.

**Figure 3.**
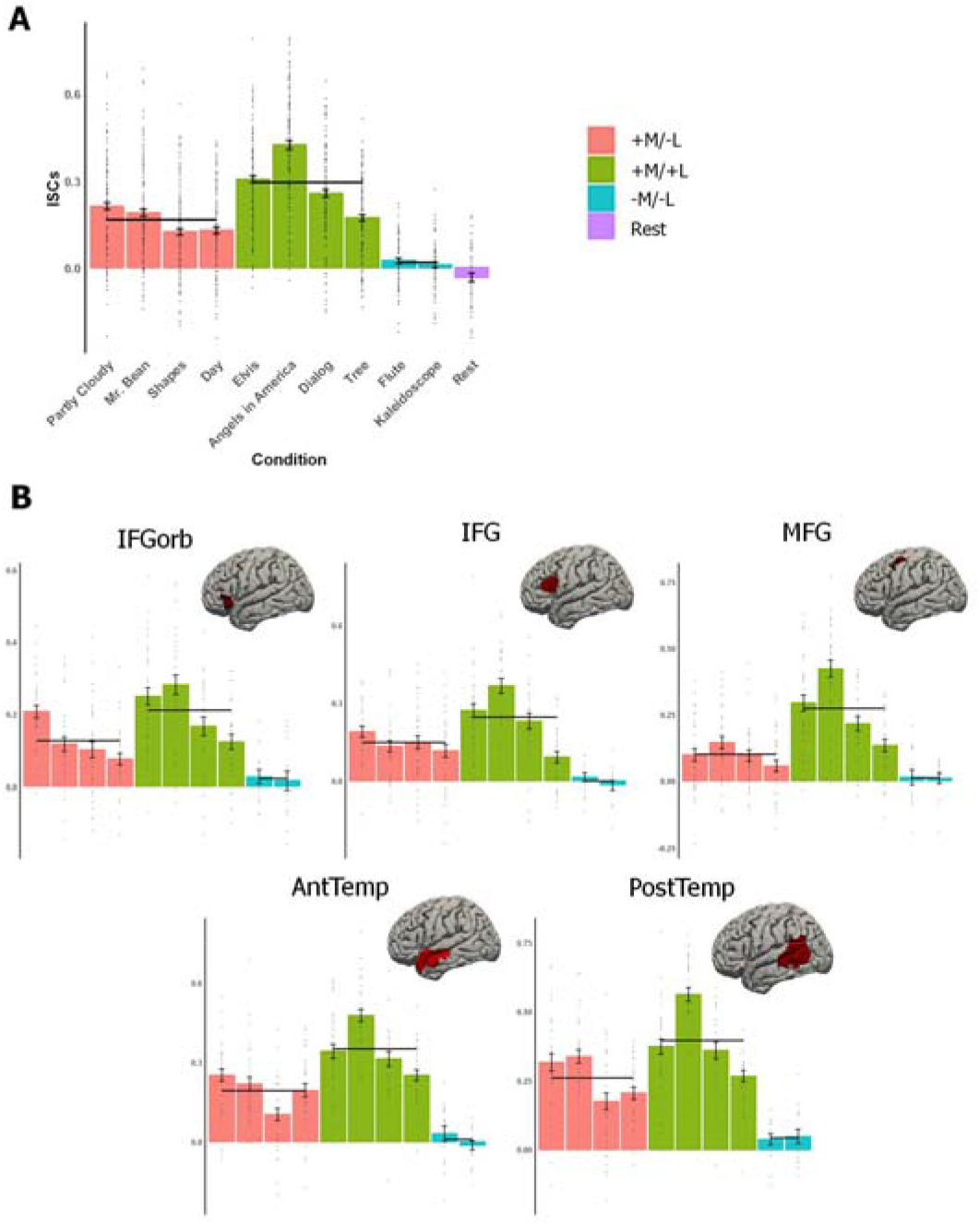
ISCs in the language network. **A.** Average ISCs across the five language fROIs. Dots indicate individual participants’ data points, and error bars indicate standard error of mean by participant. Black horizontal lines indicate the mean ISC values for each of the three condition types. **B.** Average ISCs were computed for each fROI separately (on the brains, we show the parcels used to define the fROIs; individual fROIs comprise 10% of each parcel, as described in the text). IFGorb = inferior frontal gyrus, orbital portion; IFG = inferior frontal gyrus; MFG = middle frontal gyrus; AntTemp = anterior temporal cortex; PostTemp = posterior temporal cortex.

To characterize this effect in greater detail, we next tested individual +M/-L conditions against the zero baseline. Each of the four +M/-L conditions elicited above-baseline ISCs (**Table 3, top**). Interestingly, the language fROIs tracked these non-verbal stimuli regardless of whether they were visual (Partly Cloudy, Mr. Bean, Shapes) or auditory, made up of sound effects (Day).

**Table 3.**
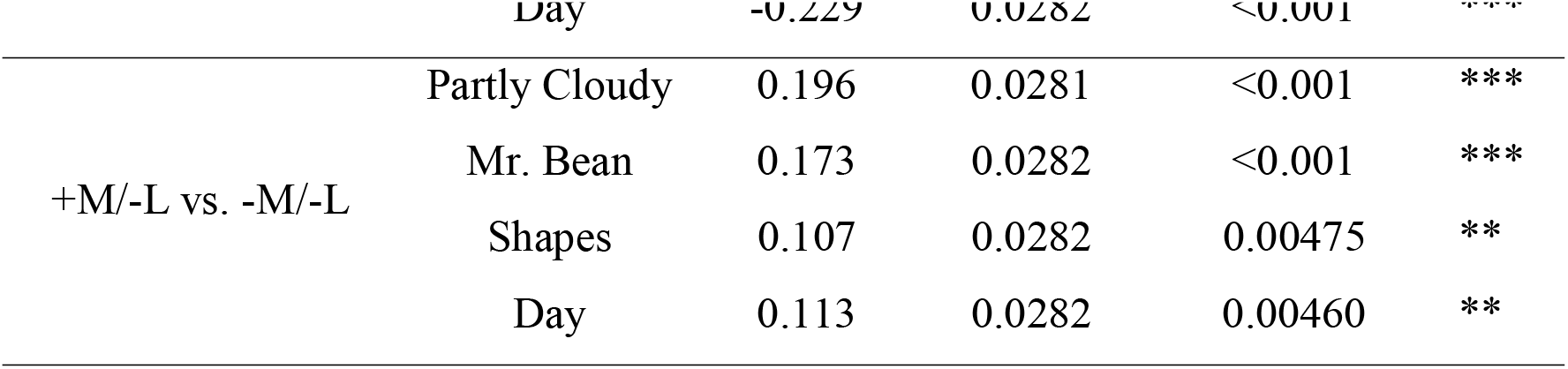
LME model statistics for the +M/-L conditions. For each condition (stimulus), ISCs were compared against the zero baseline, average +M/+L ISCs, and average -M/-L ISCs using LME models with condition as a fixed effect and participant and fROI as random intercepts. For each analysis, the following three values are reported: beta, standard error of the mean (SE), p-value (FDR-corrected for multiple comparison across baselines (n=3)). For p-values, *<0.05, **<0.01, ***<0.001. +M/-L vs. -M/-L

We next asked whether the recruitment of the language network by +M/-L stimuli can be observed at the level of individual fROIs (**Figure 3B**). Indeed, these conditions elicited ISCs that were significantly higher than the zero baseline in each of the five fROIs (**Table 4**). These results also held for each condition individually, except for Day in MFG (**Table A.1**).

**Table 4.** LME model statistics for each fROI. For each fROI, ISCs were compared across conditions with condition type as a fixed effect and participant and condition as random intercepts. +M/-L condition type was used as reference. For each fROI, the following three values are reported: beta, standard error of mean (SE), p-value (FDR-corrected for multiple comparison across fROIs (n=5)). IFGorb = inferior frontal gyrus, orbital portion; IFG = inferior frontal gyrus; MFG = middle frontal gyrus; AntTemp = anterior temporal lobe; PostTemp = posterior temporal lobe. For p-values, *<0.05, **<0.01, ***<0.001.

| fROI | beta | SE | p-value |  |
| --- | --- | --- | --- | --- |
| IFGorb | 0.127 | 0.0278 | 0.00126 | ** |
| IFG | 0.149 | 0.0353 | 0.00186 | ** |
| MFG | 0.0976 | 0.0380 | 0.0260 | * |
| AntTemp | 0.192 | 0.0337 | <0.001 | *** |
| PostTemp | 0.262 | 0.0428 | <0.001 | *** |

Taken together, these analyses show that naturalistic meaningful non-verbal stimuli, including both visual and auditory ones, elicit reliable stimulus-related activity across the language network, as indexed by reliable ISCs.

### 3.2. The language regions track meaningful non-verbal stimuli more reliably than non-meaningful stimuli, but less reliably than verbal stimuli

We next asked how the ISCs elicited by the +M/-L conditions differ from the ISCs elicited by the other two condition types. We found that the +M/-L conditions elicited ISCs (mean ISCs = 0.193, beta = 0.168, SE = 0.0434, p = 0.00151) that were higher than those elicited by non-meaningful non-verbal (-M/-L) conditions (mean ISCs = 0.00795, beta = 0.146, SE = 0.0597, p = 0.0350) but lower than those elicited by meaningful linguistic (+M/+L) conditions (mean ISCs = 0.350, beta = 0.118, SE = 0.0483, p = 0.0353). This pattern also held for each individual condition (**Table 3, middle, bottom**), and when each fROI was analyzed separately (**Table A.2**; the differences among the language fROIs in the exact patterns of ISCs are likely due to differences among the regions with respect to their sensitivity to stimulus dimensions other than the presence of semantic / verbal content). Taken together, these analyses show that meaningful non-verbal stimuli elicit stimulus-related activity across the language network that is stronger than that elicited by non-meaningful non-verbal stimuli (e.g., a musical piece) but weaker than that elicited by meaningful verbal stimuli (e.g., a story).

### 3.3. Elevated ISCs in the +L conditions cannot be explained by the amount of semantic content

Does the difference in ISCs elicited by the +M/+L versus the +M/-L conditions reflect a difference in semantic richness? To test this idea, we quantified the degree of semantic richness for each stimulus using the content unit (CU) metric—the number of referenced concepts in verbal descriptions of the stimuli (**Figure 4A**; Agis et al., 2016; Gallée et al., 2021). As a sanity check, we first examined the CU counts for the -M conditions. Because participants are prompted to provide a verbal description for each portion of the stimulus, the CU count does not go to zero for any given stimulus. Nonetheless, the CU count is significantly lower in the -M/-L conditions compared to both the +M/-L (beta = 6.48, SE = 2.19, p = 0.0143) and the +M/+L conditions (beta = 5.68, SE = 2.18, p = 0.0267), in line with our classification of +M versus -M stimuli.

**Figure 4.**
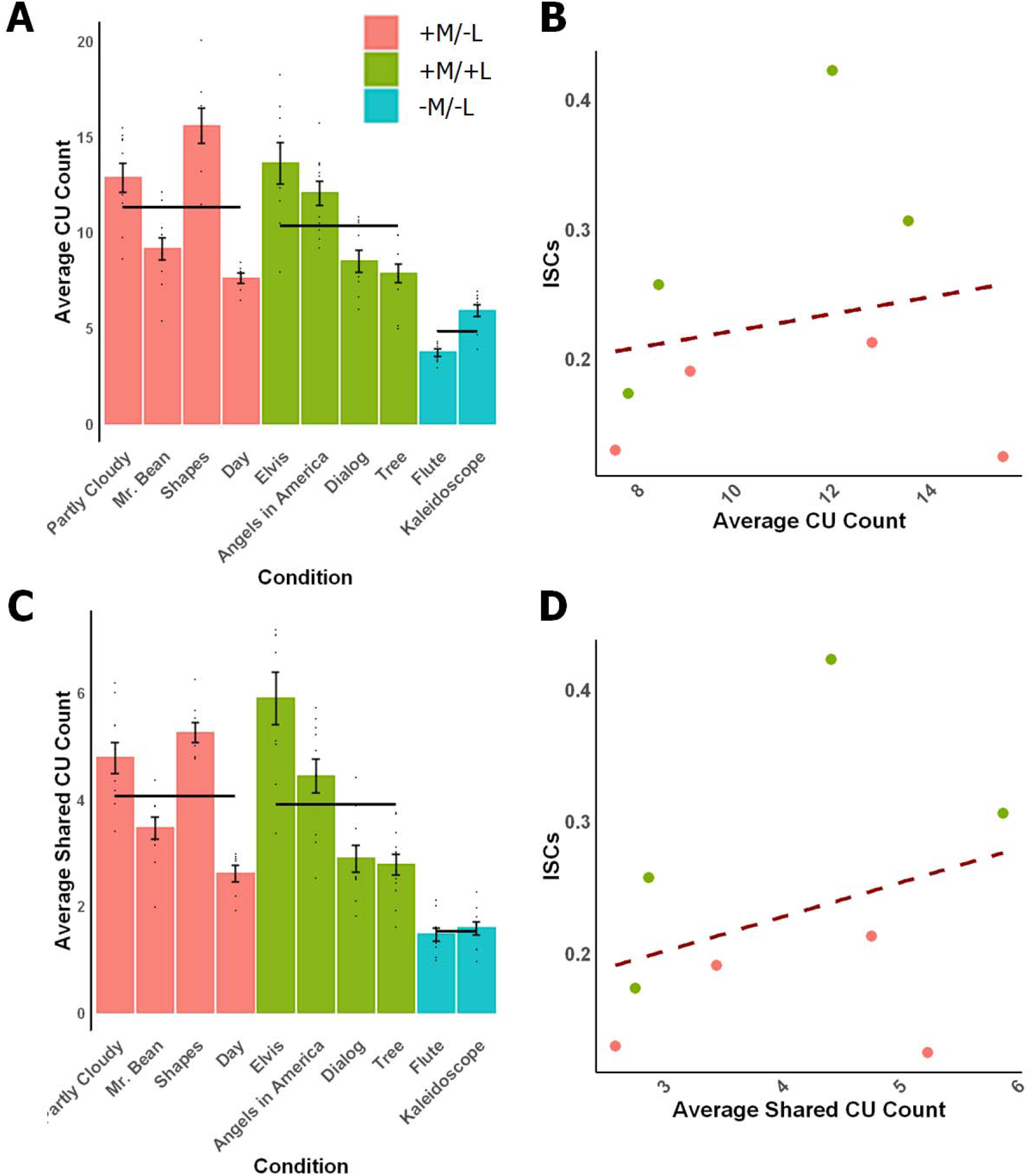
CU count cannot explain the elevated ISCs in +L stimuli. **A.** Average CU count across participants. Dots indicate individual subclips that comprise each stimulus, and error bars indicate standard error of the mean by subclip. Black horizontal lines indicate the mean CU counts for each of the three condition types. **B.** The correlation between the average CU count and the ISC across the +M stimuli. Dashed line indicates best fit. line. **C, D**. Same as A and B, but using average shared CU count.

We then tested whether the +M/+L conditions were more semantically rich compared to the +M/-L conditions. These two conditions did not differ statistically (beta = -0.800, SE = 1.96, p = 0.694; +M/-L as reference level). Furthermore, the average CU count did not reliably correlate with ISC strength for the conditions in these two condition types (**Figure 4B**; r = 0.196, p = 0.642). To test this effect more rigorously, we created two models that use CU count as a predictor of ISC strength. In the first model, we added normalized CU count as fixed effect on top of the condition type. In this model, the CU count did not have a significant effect (beta = 0.0101, SE = 0.00785, p = 0.227), and the likelihood ratio test showed that adding this extra fixed effect did not significantly improve the model (X^2^(1) = 1.54, p = 0.214). In the second model, we replaced the fixed effect: from condition type to normalized CU count. The normalized CU count had significant explanatory power in the ISCs (beta = 0.0196, SE = 0.00869, p = 0.0479), in line with the low CU count for -M stimuli. Nonetheless, the model with condition type as the fixed effect had a lower AIC (ΔAIC = -4.79), suggesting that condition type explains the ISCs better than the CU count.

To assess the generalizability of this result across metrics, we performed the same analysis using average pairwise shared CU count, which measures the similarity in the verbal description across participants and thus parallels (conceptually) the ISC measure for BOLD signals. This analysis also revealed no significant difference in the shared CU count between the two condition types (beta = -0.0348, SE = 0.823, p = 0.967; +M/-L as reference level) and no significant correlation between the average shared CU count and the ISCs (**Figures 4C, D**; r = 0.324, p = 0.433). Next, we tested the effect of shared CU count using the two LME models described above. Adding a normalized shared CU count as a fixed effect did not have a significant effect (beta = 0.0295, SE = 0.0184, p = 0.141), and the likelihood ratio test showed that adding this extra fixed effect did not significantly improve the model (X^2^(1) =2.29, p = 0.130). In the second model, the normalized shared CU count had significant explanatory power in the ISCs (beta = 0.0541, SE = 0.197, p = 0.0210) but the model with condition type as the fixed effect had a lower AIC (ΔAIC = -3.31). Altogether, these analyses rule out the interpretation that the elevated ISCs in +L conditions result from these stimuli being semantically richer.

### 3.4. The language network’s tracking of meaningful non-verbal stimuli cannot be explained by the peak timepoints

In the final analysis, we asked whether the relatively high ISCs in the language network for meaningful non-verbal conditions could be explained by a small number of peak timepoints (when language processing may be likely to be engaged). We recomputed the ISCs after removing peak (and, for comparison, trough) time points (**Figure 2C**). The peak-/trough-removed ISCs were compared against ISCs computed after random time points were removed (matched in number to the peak time points removed) and to each other (**Figure 5**). The peak-removed ISCs were significantly lower than random-timepoints-removed ISCs for each of the four +M/-L conditions (ps < 0.001, FDR corrected (n = 4)). The trough-removed ISCs were also lower than random-timepoints-removed ISCs (ps < 0.001, FDR corrected). However, although numerically, peak removal led to a larger reduction in the strength of ISCs than trough removal across conditions, this difference was only significant in one of the four conditions (Mr. Bean, t(338) = -3.68, p = 0.00108; FDR-corrected). These results suggest that although in some cases occasional strong responses in the language regions during the processing of meaningful non-verbal stimuli (perhaps related to activating linguistic representations) may contribute to the strength of ISCs for those conditions, these peak points cannot fully explain reliable tracking of these stimuli. In fact, in the +M/-L condition that elicited the highest ISCs in the language regions (Partly Cloudy), peak removal and trough removal resulted in a similar reduction in ISC strength.

**Figure 5.**
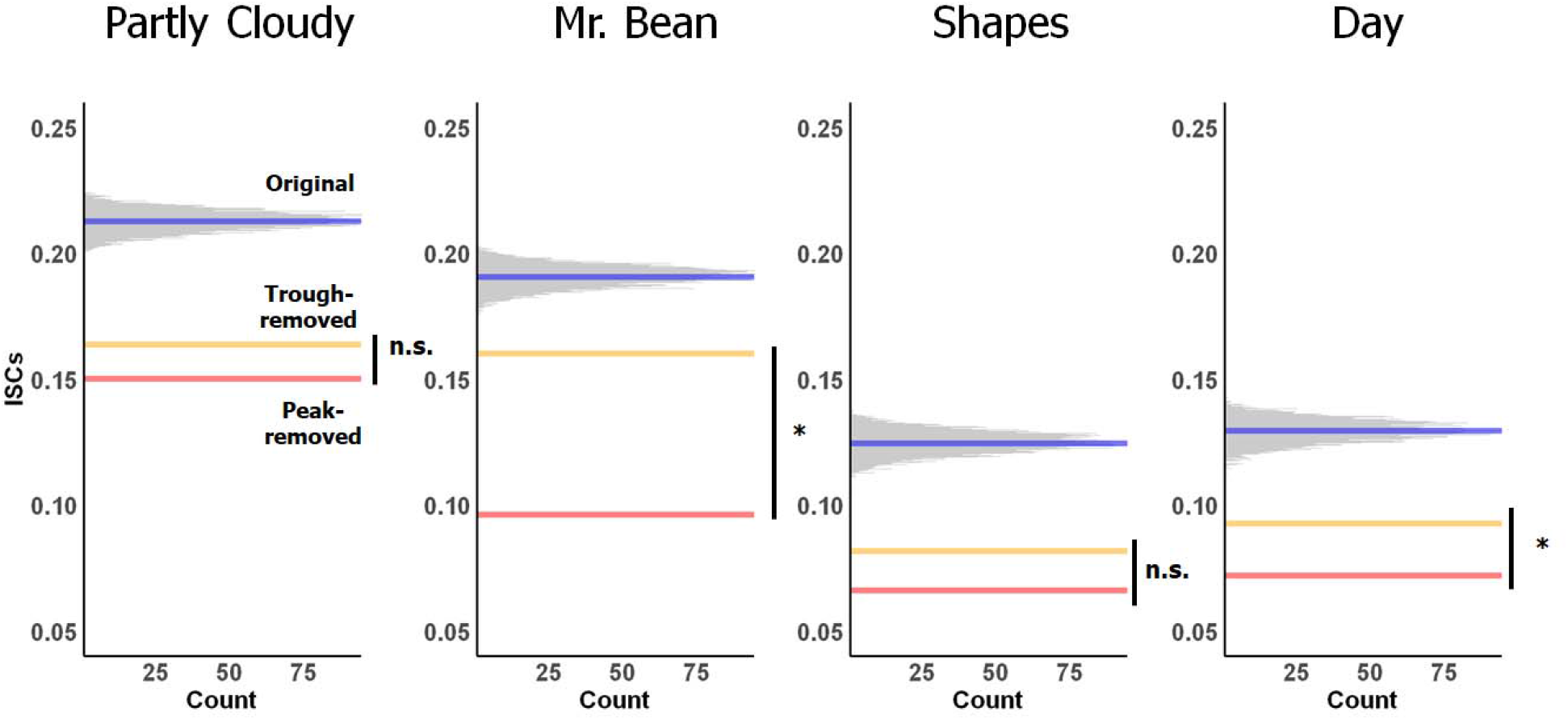
Peak-removal analysis. Peak- / trough-removed ISCs were compared against random-timepoints-removed ISCs and against each other for each +M/-L condition. The histograms show the random-timepoints-removed ISCs (generated 10,000 times; averaged across fROIs/participants) with bin width of 0.0001. The blue lines correspond to the true ISCs, the red lines – to the peak-removed ISCs, and the yellow lines – to the trough-removed ISCs. For p-values, *<0.05, n.s. = non-significant (p > 0.05).

## 4. Discussion

A left-lateralized network of frontal and temporal areas has long been implicated in language comprehension. However, the role of the language network in non-verbal semantic processing remains debated. Using the inter-subject correlation (ISC) approach (Hasson et al., 2008), we examined the degree of ‘tracking’ of meaningful (i.e., semantically rich) non-verbal stimuli by language areas, identified functionally in each individual (e.g., Fedorenko et al., 2010). We found that i) the language regions reliably track meaningful non-verbal stimuli (both visual and auditory); ii) the degree of tracking of such stimuli is higher than for non-meaningful stimuli (like music or dynamically-changing kaleidoscope images) but lower than for verbal stimuli, in spite of the verbal and non-verbal meaningful stimuli being similarly semantically rich; and iii) the tracking of meaningful non-verbal stimuli cannot be explained by the presence of occasional scenes that elicit a strong response in the language regions (e.g., due to activating some linguistic representations).

The strong tracking of naturalistic verbal stimuli by the language network replicates several past findings (e.g., Wilson et al., 2007; Lerner et al., 2011; Blank & Fedorenko, 2017, 2020). Importantly, our analysis of content units (Agis et al., 2016; Gallée et al., 2021) demonstrated that such elevated tracking (relative to non-verbal meaningful stimuli) is not merely a result of verbal stimuli containing higher semantic content That said, we acknowledge that verbal and non-verbal meaningful stimuli (in general, or for the particular set used in the current study) may differ under other metrics of semantic content. The reliably stronger tracking of verbal than non-verbal stimuli in the language network adds to the growing body of evidence suggesting that the language network is highly selective for language processing relative to diverse non-linguistic functions (e.g., Monti et al., 2009; Fedorenko et al., 2011; Pritchett et al., 2018; Ivanova et al., 2020; Paunov et al., 2022).

The observed difference between the verbal and non-verbal meaningful stimuli requires care in its interpretation. Lower ISCs in the +M/-L compared to the +M/+L conditions imply *lower consistency* in responses to these stimuli across participants, but not necessarily lower evoked responses. In other words, this result on its own does not rule out the possibility that the language network is similarly responsive to linguistic and non-linguistic meaningful content. However, previous work argues against this possibility. Specifically, Ivanova et al. (2021) found that event pictures activate the language network to a lesser degree compared to semantically-matched sentences (discussed in more detail below). A similar result was obtained using auditory meaningful stimuli (Humphries et al., 2001). These finding suggest that the more likely explanation for the lower ISCs for the meaningful non-verbal (+M/-L) stimuli is the language network’s lower degree of engagement with those stimuli rather than merely a lower degree of response consistency.

Worth a special mention is a very low (near-zero) degree of tracking by the language network of a naturalistic musical stimulus. Although a number of neuroimaging studies have now provided evidence against the role of the language network in music perception, including in the processing of music structure (e.g., Fedorenko et al., 2011; Rogalsky et al., 2011; Deen et al., 2015; Chen et al., 2023), many researchers continue to argue for overlapping mechanisms between language and music (e.g., Sammler et al., 2013; Kunert et al., 2015). The lack of stimulus-related activity in the language network (including its inferior frontal components (‘Broca’s area’)) during naturalistic listening of a musical piece provides compelling evidence that the language areas are not performing computations related to the musical stimulus, including its structure (the most common claim; e.g., Kunert et al., 2015). Future research using intra-subject correlations based on repeated exposure to the same music stimulus can further clarify whether this result is due to the lack of engagement of the language network during music listening or due to the interpretive freedom associated with music stimuli.

Our main finding—above-baseline tracking by the language network of meaningful non-verbal stimuli—is in line with recent work showing that a) event pictures elicit a reliable response in the language network, but b) this response is lower than that elicited by semantically matched sentences (Ivanova et al., 2021). Here, we extend these results in an important way by ruling out the possibility that the responses in the language network to visual events are task-driven. In particular, it has been suggested that when viewing visual objects and events, participants may activate words related to those stimuli in order to facilitate task performance (Trueswell & Papafragou, 2010; Greene & Fei-Fei, 2014). However, naturalistic stimuli presented under passive-viewing instructions are unlikely to lead to verbal re-coding given no obvious advantages of such re-coding. And the possibility of occasional verbal re-coding—perhaps in the parts of the stimuli that impose high processing or memory demands—is further ruled out by our peak-removal analysis, which showed that the observed ISCs are not driven by a small number of time points with elevated activity in the language network.

Together, our results suggest that the language network exhibits continuous and automatic tracking of semantic content in non-verbal stimuli. What does this tracking reflect? Data from neuropsychological investigations of patients with acquired brain damage (e.g., Whitehouse et al., 1978; Chertkow et al., 1997; Saygin et al., 2004; Colvin et al., 2019; Ivanova et al., 2021; Warren & Dickey, 2021) demonstrate that damage to the language network does not impair visual / abstract semantic processing. For example, Ivanova et al. (2021; see also Dickey & Warren, 2015) showed that some individuals with aphasia (including severe aphasia) perform well on a challenging semantic task on event pictures (judging event plausibility for events with two animate participants; e.g., a jester entertaining a king vs. a king entertaining a jester). However, it is still possible that the language network contributes to semantic processing— redundantly with other systems—without being crucial for it. Another possibility, suggested by a reviewer, is that the relatively high ISCs in the language areas during the processing of meaningful non-verbal stimuli reflect the processing of communicative elements present in those stimuli, like facial expressions or body language. However, previous studies have found that the language network does not respond to facial expressions, eye gaze, or even speech-accompanying gestures (e.g., Deen et al., 2015; Pritchett et al., 2018; Jouravlev et al., 2019), which makes this possibility unlikely.

Understanding what precise semantic computations or aspects of meaning drive the language network’s response to non-verbal meaningful stimuli is an important future endeavor, which can provide important clues about the relationship between linguistic and semantic processing. One constraint on this space of hypotheses has to do with the size of the language network’s ‘temporal integration (or receptive) window’ (e.g., Lerner et al., 2011). In particular, previous work has shown that the temporal integration window of the language network is relatively short, on the order of a clause or sentence (e.g., Lerner et al., 2011; Blank & Fedorenko, 2020; Regev, Casto et al., 2022; Shain, Kean et al., 2023). It is therefore likely that non-verbal meanings that the language system is concerned with have to do with individually described states or agent-patient interactions, but not longer-timescale representations like situation models whose construction can span long sequences of events (Johnson-Laird, 1983; Zwaan & Radvansky, 1998; Loschky et al., 2019). But whether the language network has a preference for particular kinds or aspects of semantic content remains to be determined. Future work that manipulates in more controlled ways various properties of visual or auditory events may help uncover such preferences, including potential dissociations among the different components of the language network.

## 5. Conclusions

Our study provides evidence that the language network reliably tracks non-verbal meaningful stimuli. The use of naturalistic stimuli and control analyses allowed us to rule out explanations in terms of broad differences in semantic richness and task-driven linguistic re-coding. Overall, these results demonstrate a role of the language network in naturalistic non-verbal semantic processing and set the stage for future work on the precise nature and functions of its contributions.

## Acknowledgments

We would like to acknowledge the Athinoula A. Martinos Imaging Center at the McGovern Institute for Brain Research at MIT, and its support team (Steve Shannon and Atsushi Takahashi). We thank former and current EvLab members (especially Olessia Jouravlev, Zach Mineroff, Brianna Pritchett, and Caitlyn Hoeflin) for help with data collection, Amaya Arcelus for help with creating the Heider & Simmel style animation, and Jeanne Gallée and Caroline Arnold for help with recording the dialog.

## Appendix A: Supplementary Data

**Table A.1.**
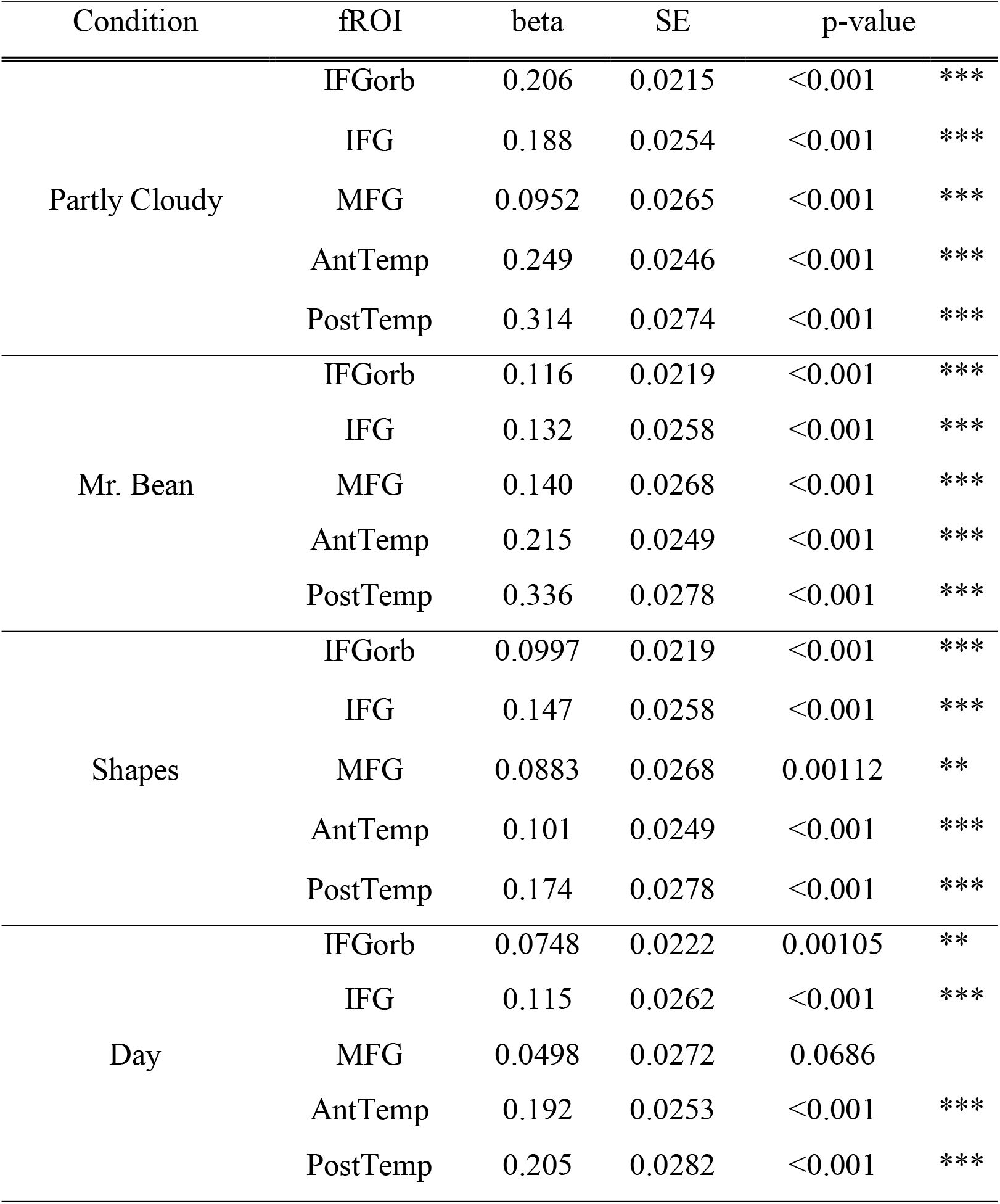
LME model statistics for each condition-fROI pair. For each fROI, ISCs were compared across conditions/fROIs with condition type as a fixed effect and participant as random intercepts (Here, each +M/-L condition are recoded into individual condition types). Zero-baseline was used as a reference. For each fROI, the following three values are reported: beta, standard error of mean (SE), p-value (FDR-corrected for multiple comparison across fROIs (n=5)). IFGorb = inferior frontal gyrus, orbital portion; IFG = inferior frontal gyrus; MFG = middle frontal gyrus; AntTemp = anterior temporal lobe; PostTemp = posterior temporal lobe. For p-values, *<0.05, **<0.01, ***<0.001.

**Table A.2.**
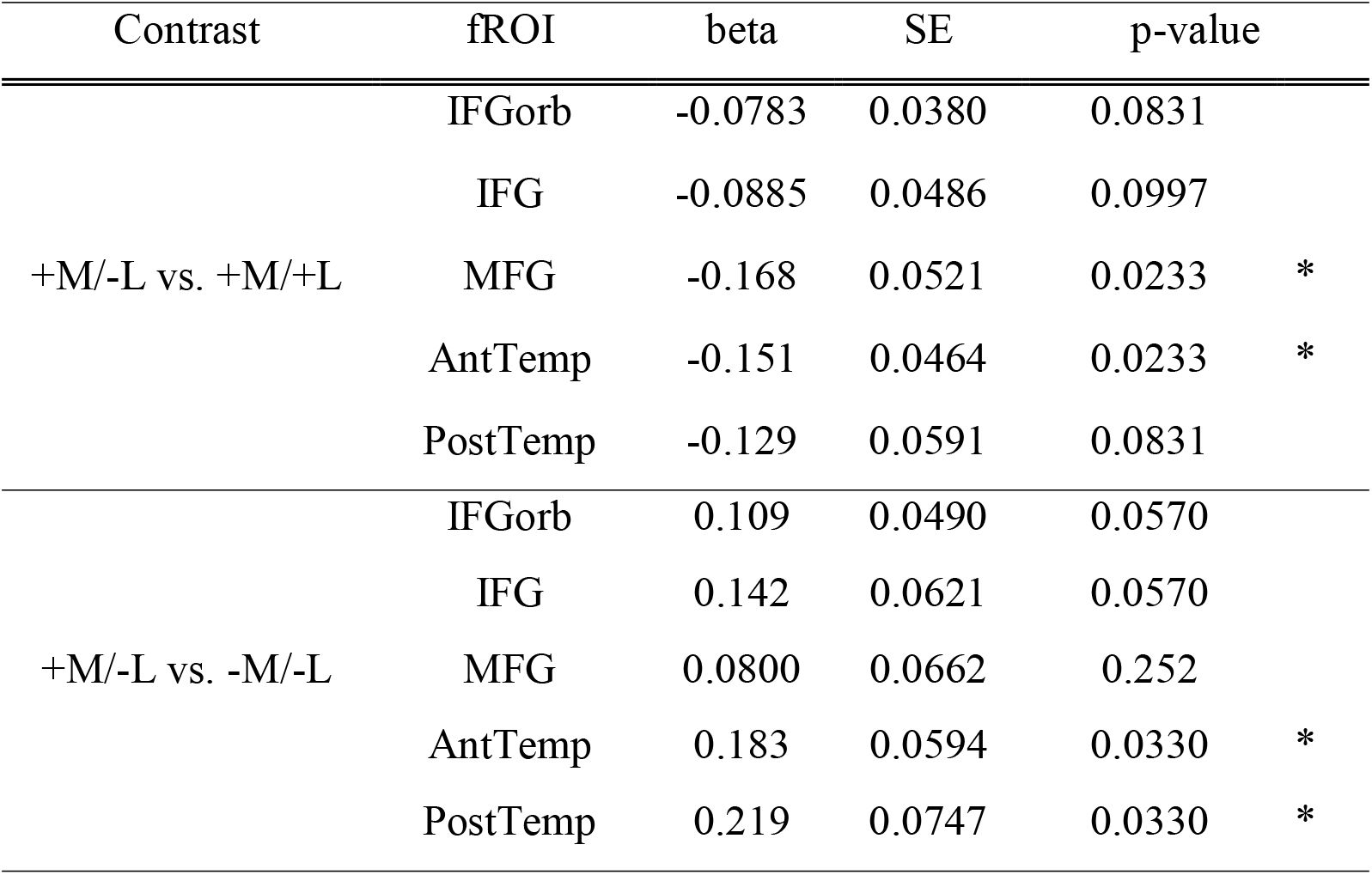
LME model statistics for each fROI on condition type comparison. For each fROI, ISCs were compared across fROIs with condition type as a fixed effect and participant and condition as random intercepts. For each fROI, the following three values are reported: beta, standard error of mean (SE), p-value (FDR-corrected for multiple comparison across fROIs (n=5)). IFGorb = inferior frontal gyrus, orbital portion; IFG = inferior frontal gyrus; MFG = middle frontal gyrus; AntTemp = anterior temporal lobe; PostTemp = posterior temporal lobe. For p-values, *<0.05.

## References

Agis, D., Goggins, M. B., Oishi, K., Oishi, K., Davis, C., Wright, A., … & Hillis, A. E. (2016). Picturing the size and site of stroke with an expanded National Institutes of Health Stroke Scale. Stroke, 47(6), 1459–1465.

Amalric, M., & Dehaene, S. (2019). A distinct cortical network for mathematical knowledge in the human brain. NeuroImage, 189, 19–31.

Amunts, K., Schleicher, A., Bürgel, U., Mohlberg, H., Uylings, H. B., & Zilles, K. (1999). Broca’s region revisited: Cytoarchitecture and intersubject variability. Journal of Comparative Neurology, 412(2), 319–341.

Apperly, I. A., Samson, D., Carroll, N., Hussain, S., & Humphreys, G. (2006). Intact first-and second-order false belief reasoning in a patient with severely impaired grammar. Social Neuroscience, 1(3–4), 334–348.

Baldassano, C., Hasson, U., & Norman, K. A. (2018). Representation of Real-World Event Schemas during Narrative Perception. Journal of Neuroscience, 38(45), 9689–9699. https://doi.org/10.1523/JNEUROSCI.0251-18.2018

Barr, D. J., Levy, R., Scheepers, C., & Tily, H. J. (2013). Random effects structure for confirmatory hypothesis testing: Keep it maximal. Journal of Memory and Language, 68(3), 255–278. https://doi.org/10.1016/j.jml.2012.11.001

Bek, J., Blades, M., Siegal, M., & Varley, R. (2010). Language and spatial reorientation: Evidence from severe aphasia. Journal of Experimental Psychology: Learning, Memory, and Cognition, 36(3), 646.

Benjamini, Y., & Yekutieli, D. (2001). The Control of the False Discovery Rate in Multiple Testing under Dependency. The Annals of Statistics, 29(4), 1165–1188.

Benn, Y., Ivanova, A. A., Clark, O., Mineroff, Z., Seikus, C., Silva, J. S., Varley, R., & Fedorenko, E. (2023). No evidence for a special role of language in feature-based categorization. Cerebral Cortex. In press

Blank, I. A., & Fedorenko, E. (2017). Domain-general brain regions do not track linguistic input as closely as language-selective regions. Journal of Neuroscience, 37(41), 9999–10011.

Braga, R. M., DiNicola, L. M., Becker, H. C., & Buckner, R. L. (2020). Situating the left-lateralized language network in the broader organization of multiple specialized large-scale distributed networks. Journal of Neurophysiology.

Brett, M., Johnsrude, I. S., & Owen, A. M. (2002). The problem of functional localization in the human brain. Nature Reviews Neuroscience, 3(3), 243–249. https://doi.org/10.1038/nrn756

Chen, X., Affourtit, J., Ryskin, R., Regev, T. I., Norman-Haignere, S., Jouravlev, O., Malik-Moraleda, S., Kean, H., Varley, R., & Fedorenko, E. (2023). The human language system, including its inferior frontal component in ’Broca’s area’, does not support music processing. Cerebral Cortex

Chertkow, H., Bub, D., Deaudon, C., & Whitehead, V. (1997). On the Status of Object Concepts in Aphasia. Brain and Language, 58(2), 203–232. https://doi.org/10.1006/brln.1997.1771

Colvin, M., Warren, T., & Dickey, M. W. (2019). Event Knowledge and Verb Knowledge Predict Sensitivity to Different Aspects of Semantic Anomalies in Aphasia. In K. Carlson, Jr. Clifton Charles, & J. D. Fodor (Eds.), Grammatical Approaches to Language Processing: Essays in Honor of Lyn Frazier (pp. 241–259). Springer International Publishing. https://doi.org/10.1007/978-3-030-01563-3_13

Deen, B., Koldewyn, K., Kanwisher, N., & Saxe, R. (2015). Functional organization of social perception and cognition in the superior temporal sulcus. Cerebral Cortex, 25(11), 4596– 4609.

Devereux, B. J., Clarke, A., Marouchos, A., & Tyler, L. K. (2013). Representational similarity analysis reveals commonalities and differences in the semantic processing of words and objects. Journal of Neuroscience, 33(48), 18906–18916.

Dickey, M. W., & Warren, T. (2015). The influence of event-related knowledge on verb-argument processing in aphasia. Neuropsychologia, 67, 63–81.

Fedorenko, E. (2021). The early origins and the growing popularity of the individual-subject analytic approach in human neuroscience. Current Opinion in Behavioral Sciences, 40, 105–112. https://doi.org/10.1016/j.cobeha.2021.02.023

Fedorenko, E., Behr, M. K., & Kanwisher, N. (2011). Functional specificity for high-level linguistic processing in the human brain. Proceedings of the National Academy of Sciences, 108(39), 16428–16433.

Fedorenko, E., & Blank, I. A. (2020). Broca’s area is not a natural kind. Trends in Cognitive Sciences.

Fedorenko, E., Hsieh, P.-J., Nieto-Castañón, A., Whitfield-Gabrieli, S., & Kanwisher, N. (2010). New method for fMRI investigations of language: Defining ROIs functionally in individual subjects. Journal of Neurophysiology, 104(2), 1177–1194.

Fedorenko, E., & Varley, R. (2016). Language and thought are not the same thing: Evidence from neuroimaging and neurological patients. Annals of the New York Academy of Sciences, 1369(1), 132.

Futrell, R., Gibson, E., Tily, H. J., Blank, I., Vishnevetsky, A., Piantadosi, S. T., & Fedorenko, E. (2021). The Natural Stories corpus: A reading-time corpus of English texts containing rare syntactic constructions. Language Resources and Evaluation, 55(1), 63–77. https://doi.org/10.1007/s10579-020-09503-7

Gallée, J., Cordella, C., Fedorenko, E., Hochberg, D., Touroutoglou, A., Quimby, M., & Dickerson, B. C. (2021). Breakdowns in informativeness of naturalistic speech production in primary progressive aphasia. Brain sciences, 11(2), 130.

Goldberg, A. E. (2010). Verbs, frames, and constructions. In M. Rappaport Hovav, E. Doron, & I. Sichel (Eds.), Syntax, lexical semantics, and event structure (pp. 39–58). Oxford University Press.

Greene, M. R., & Fei-Fei, L. (2014). Visual categorization is automatic and obligatory: Evidence from Stroop-like paradigm. Journal of Vision, 14(1), 14. https://doi.org/10.1167/14.1.14

Handjaras, G., Leo, A., Cecchetti, L., Papale, P., Lenci, A., Marotta, G., Pietrini, P., & Ricciardi, E. (2017). Modality-independent encoding of individual concepts in the left parietal cortex. Neuropsychologia, 105, 39–49.

Hasson, U., Egidi, G., Marelli, M., & Willems, R. M. (2018). Grounding the neurobiology of language in first principles: The necessity of non-language-centric explanations for language comprehension. Cognition, 180, 135–157.

Hasson, U., Furman, O., Clark, D., Dudai, Y., & Davachi, L. (2008). Enhanced Intersubject Correlations during Movie Viewing Correlate with Successful Episodic Encoding. Neuron, 57(3), 452–462. https://doi.org/10.1016/j.neuron.2007.12.009

Hasson, U., Nir, Y., Levy, I., Fuhrmann, G., & Malach, R. (2004). Intersubject synchronization of cortical activity during natural vision. Science, 303(5664), 1634–1640.

HBO (Producer). (2003, December 7). Chapter 1: Bad News [Audio only]. Angels in America. Retrieved from https://www.imdb.com/title/tt0318997/

Heider, F., & Simmel, M. (1944). An experimental study of apparent behavior. The American Journal of Psychology, 57(2), 243–259.

Humphries, C., Willard, K., Buchsbaum, B., & Hickok, G. (2001). Role of anterior temporal cortex in auditory sentence comprehension: an fMRI study. Neuroreport, 12(8), 1749–1752.

Ivanova, A. A., Mineroff, Z., Zimmerer, V., Kanwisher, N., Varley, R., & Fedorenko, E. (2021). The Language Network Is Recruited but Not Required for Nonverbal Event Semantics. Neurobiology of Language, 2(2), 176–201. https://doi.org/10.1162/nol_a_00030

Ivanova, A. A., Srikant, S., Sueoka, Y., Kean, H. H., Dhamala, R., O’Reilly, U.-M., Bers, M. U., & Fedorenko, E. (2020). Comprehension of computer code relies primarily on domain-general executive brain regions. ELife, 9, e58906. https://doi.org/10.7554/eLife.58906

Johnson-Laird, P. N. (1983). Mental Models: Towards a Cognitive Science of Language, Inference, and Consciousness. Harvard University Press.

Jouravlev, O., Zheng, D., Balewski, Z., Pongos, A. L. A., Levan, Z., Goldin-Meadow, S., & Fedorenko, E. (2019). Speech-accompanying gestures are not processed by the language-processing mechanisms. Neuropsychologia, 132, 107132.

Katz, J. J., & Fodor, J. A. (1963). The structure of a semantic theory. language, 39(2), 170–210.

Kunert, R., Willems, R. M., Casasanto, D., Patel, A. D., & Hagoort, P. (2015). Music and Language Syntax Interact in Broca’s Area: An fMRI Study. PLOS ONE, 10(11), e0141069. https://doi.org/10.1371/journal.pone.0141069

Lambon Ralph, M. A., Jefferies, E., Patterson, K., & Rogers, T. T. (2017). The neural and computational bases of semantic cognition. Nature Reviews Neuroscience, 18(1), 42–55.

Lecours, A., & Joanette, Y. (1980). Linguistic and other psychological aspects of paroxysmal aphasia. Brain and Language, 10(1), 1–23. https://doi.org/10.1016/0093-934X(80)90034-6

Lerner, Y., Honey, C. J., Silbert, L. J., & Hasson, U. (2011). Topographic mapping of a hierarchy of temporal receptive windows using a narrated story. Journal of Neuroscience, 31(8), 2906– 2915.

Lipkin, B., Affourtit, J., Small, H., Mineroff, Z., Nieto-Castañón, A., and Fedorenko, E. (in prep.). In defense of individual-level functional neural markers: Evidence from large-scale fMRI datasets of functional ‘localizers’ for the language and the Multiple Demand networks.

Lipkin, B., Tuckute, G., Affourtit, J., Small, H., Mineroff, Z., Kean, H., Jouravlev, O., Rakocevic, L., Pritchett, B., Siegelman, M., Hoeflin, C., Pongos, A., Blank, I. A., Struhl, M. K., Ivanova, A., Shannon, S., Sathe, A., Hoffmann, M., Nieto-Castañón, A., & Fedorenko, E. (2022). LanA (Language Atlas): A probabilistic atlas for the language network based on fMRI data from >800 individuals (p. 2022.03.06.483177). bioRxiv. https://doi.org/10.1101/2022.03.06.483177

Liu, Y.-F., Kim, J., Wilson, C., & Bedny, M. (2020). Computer code comprehension shares neural resources with formal logical inference in the fronto-parietal network. ELife, 9, e59340. https://doi.org/10.7554/eLife.59340

Loschky, L. C., Larson, A. M., Smith, T. J., & Magliano, J. P. (2020). The Scene Perception & Event Comprehension Theory (SPECT) Applied to Visual Narratives. Topics in Cognitive Science, 12(1), 311–351. https://doi.org/10.1111/tops.12455

Mahowald, K., & Fedorenko, E. (2016). Reliable individual-level neural markers of high-level language processing: A necessary precursor for relating neural variability to behavioral and genetic variability. Neuroimage, 139, 74–93.

Mineroff, Z., Blank, I. A., Mahowald, K., & Fedorenko, E. (2018). A robust dissociation among the language, multiple demand, and default mode networks: Evidence from inter-region correlations in effect size. Neuropsychologia, 119, 501–511. https://doi.org/10.1016/j.neuropsychologia.2018.09.011

Monti, M. M., Parsons, L. M., & Osherson, D. N. (2009). The boundaries of language and thought in deductive inference. Proceedings of the National Academy of Sciences, 106(30), 12554–12559.

Monti, M. M., Parsons, L. M., & Osherson, D. N. (2012). Thought beyond language: Neural dissociation of algebra and natural language. Psychological Science, 23(8), 914–922.

Mr Bean. (2009, Aug 25). Falling Asleep in Church | Funny Clip | Mr Bean Official [Video file]. Retrieved from https://www.youtube.com/watch?v=bhg-ZZ6WA

Nieto-Castañón, A., & Fedorenko, E. (2012). Subject-specific functional localizers increase sensitivity and functional resolution of multi-subject analyses. Neuroimage, 63(3), 1646– 1669.

Oldfield, R. C. (1971). The assessment and analysis of handedness: The Edinburgh inventory. Neuropsychologia, 9(1), 97–113.

Patterson, K., Nestor, P. J., & Rogers, T. T. (2007). Where do you know what you know? The representation of semantic knowledge in the human brain. Nature reviews neuroscience, 8(12), 976–987.

Paunov, A. A. M. (2018). *FMRI studies of the relationship between language and theory of mind in adult cognition* [PhD Thesis]. Massachusetts Institute of Technology.

Paunov, A. M., Blank, I. A., Jouravlev, O., Minnerof, Z., Gallée, J., & Fedorenko, E. (2022). Differential tracking of linguistic vs. Mental state content in naturalistic stimuli by language and Theory of Mind (ToM) brain networks. Neurobiology of Language, 1–67. https://doi.org/10.1162/nol_a_00071

Pritchett, B. L., Hoeflin, C., Koldewyn, K., Dechter, E., & Fedorenko, E. (2018). High-level language processing regions are not engaged in action observation or imitation. Journal of Neurophysiology, 120(5), 2555–2570.

Pustejovsky, J. (2005) Lexical Semantics: Overview in Encyclopedia of Language and Linguistics, second edition, Volumes 1–14

Rogalsky, C., Rong, F., Saberi, K., & Hickok, G. (2011). Functional Anatomy of Language and Music Perception: Temporal and Structural Factors Investigated Using Functional Magnetic Resonance Imaging. Journal of Neuroscience, 31(10), 3843–3852. https://doi.org/10.1523/JNEUROSCI.4515-10.2011

Sammler, D., Koelsch, S., Ball, T., Brandt, A., Grigutsch, M., Huppertz, H.-J., Knösche, T. R., Wellmer, J., Widman, G., Elger, C. E., Friederici, A. D., & Schulze-Bonhage, A. (2013). Co-localizing linguistic and musical syntax with intracranial EEG. NeuroImage, 64, 134–146. https://doi.org/10.1016/j.neuroimage.2012.09.035

Saxe, R., Brett, M., & Kanwisher, N. (2006). Divide and conquer: A defense of functional localizers. NeuroImage, 30(4), 1088–1096; discussion 1097-1099. https://doi.org/10.1016/j.neuroimage.2005.12.062

Saygın, A. P., Wilson, S. M., Dronkers, N. F., & Bates, E. (2004). Action comprehension in aphasia: Linguistic and non-linguistic deficits and their lesion correlates. Neuropsychologia, 42(13), 1788–1804. https://doi.org/10.1016/j.neuropsychologia.2004.04.016

Scott, T. L., Gallée, J., & Fedorenko, E. (2017). A new fun and robust version of an fMRI localizer for the frontotemporal language system. Cognitive Neuroscience, 8(3), 167–176.

Shain, C., Kean, H., Lipkin, B., Affourtit, J., Siegelman, M., Mollica, F., & Fedorenko, E. (2023). Graded sensitivity to structure and meaning throughout the human language network. bioRxiv.

Shain, C., Paunov, A., Chen, X., Lipkin, B., & Fedorenko, E. (2022). No evidence of theory of mind reasoning in the human language network. bioRxiv, 2022-07.

Silbert, L. J., Honey, C. J., Simony, E., Poeppel, D., & Hasson, U. (2014). Coupled neural systems underlie the production and comprehension of naturalistic narrative speech. Proceedings of the National Academy of Sciences, 111(43), E4687–E4696.

Silver, N. C., & Dunlap, W. P. (1987). Averaging correlation coefficients: Should Fisher’s z transformation be used? Journal of Applied Psychology, 72(1), 146–148. https://doi.org/10.1037/0021-9010.72.1.146

Regev, T. I., Casto, C., Hosseini, E. A., Adamek, M., Brunner, P., & Fedorenko, E. (2022). Intracranial recordings reveal three distinct neural response patterns in the language network. bioRxiv, 2022-12.

Thierry, G., & Price, C. J. (2006). Dissociating Verbal and Nonverbal Conceptual Processing in the Human Brain. Journal of Cognitive Neuroscience, 18(6), 1018–1028. https://doi.org/10.1162/jocn.2006.18.6.1018

Tomaiuolo, F., MacDonald, J. D., Caramanos, Z., Posner, G., Chiavaras, M., Evans, A. C., & Petrides, M. (1999). Morphology, morphometry and probability mapping of the pars opercularis of the inferior frontal gyrus: An in vivo MRI analysis. European Journal of Neuroscience, 11(9), 3033–3046.

Trueswell, J. C., & Papafragou, A. (2010). Perceiving and remembering events cross-linguistically: Evidence from dual-task paradigms. Journal of Memory and Language, 63(1), 64–82. https://doi.org/10.1016/j.jml.2010.02.006

Vagharchakian, L., Dehaene-Lambertz, G., Pallier, C., & Dehaene, S. (2012). A temporal bottleneck in the language comprehension network. Journal of Neuroscience, 32(26), 9089–9102.

Varley, R. A., Klessinger, N. J., Romanowski, C. A., & Siegal, M. (2005). Agrammatic but numerate. Proceedings of the National Academy of Sciences, 102(9), 3519–3524.

Varley, R., & Siegal, M. (2000). Evidence for cognition without grammar from causal reasoning and ‘theory of mind’in an agrammatic aphasic patient. Current Biology, 10(12), 723–726.

Varley, R., Siegal, M., & Want, S. C. (2001). Severe impairment in grammar does not preclude theory of mind. Neurocase, 7(6), 489–493.

Visser, M., Jefferies, E., Embleton, K. V., & Lambon Ralph, M. A. (2012). Both the middle temporal gyrus and the ventral anterior temporal area are crucial for multimodal semantic processing: Distortion-corrected fMRI evidence for a double gradient of information convergence in the temporal lobes. Journal of Cognitive Neuroscience, 24(8), 1766–1778.

Warren, T., & Dickey, M. W. (2021). The use of linguistic and world knowledge in language processing. Language and Linguistics Compass, 15(4), e12411.

Warrington, E. K. (1975). The selective impairment of semantic memory. The Quarterly journal of experimental psychology, 27(4), 635–657.

Whitehouse, P., Caramazza, A., & Zurif, E. (1978). Naming in aphasia: Interacting effects of form and function. Brain and Language, 6(1), 63–74. https://doi.org/10.1016/0093-934X(78)90044-5

Whitfield-Gabrieli, S., & Nieto-Castanon, A. (2012). Conn: A functional connectivity toolbox for correlated and anticorrelated brain networks. Brain Connectivity, 2(3), 125–141.

Willems, R. M., Benn, Y., Hagoort, P., Toni, I., & Varley, R. (2011). Communicating without a functioning language system: Implications for the role of language in mentalizing. Neuropsychologia, 49(11), 3130–3135.

Willems, R. M., Van der Haegen, L., Fisher, S. E., & Francks, C. (2014). On the other hand: Including left-handers in cognitive neuroscience and neurogenetics. Nature Reviews Neuroscience, 15(3), 193–201.

Wilson, S. M., Molnar-Szakacs, I., & Iacoboni, M. (2007). Beyond superior temporal cortex: Intersubject correlations in narrative speech comprehension. Cerebral Cortex, 18(1), 230– 242.

Wurm, M. F., & Caramazza, A. (2019). Distinct roles of temporal and frontoparietal cortex in representing actions across vision and language. Nature Communications, 10(1), 289. https://doi.org/10.1038/s41467-018-08084-y

Zwaan, R. A., & Radvansky, G. A. (1998). Situation models in language comprehension and memory. Psychological Bulletin, 123(2), 162–185. https://doi.org/10.1037/0033-2909.123.2.162

